# In-vivo optogenetic manipulation approach for gerbil medial nucleus of trapezoid body

**DOI:** 10.1101/2025.03.10.642510

**Authors:** Ben-Zheng Li, Matthew Ridenour, Shani Poleg, Praveen Kuruppath, Elizabeth A. McCullagh, Tim Lei, Achim Klug

## Abstract

**Purpose:** This study introduces an in-vivo optogenetic manipulation approach for the medial nucleus of the trapezoid body (MNTB) in Mongolian gerbils, a species with a hearing range similar to humans. The MNTB is crucial for sound localization, but traditional methods lack temporal precision and reversibility. The aim of this study is to develop a specific, reversible method for controlling MNTB activity with fast and precise temporal control, an approach vital for studying sound localization.

**Methods:** We stereotactically injected adeno-associated viral vectors encoding opsins into the gerbil MNTB. Precise targeting was achieved despite the MNTB’s location in a deep, heavily myelinated area of the brainstem. Opsin expression was confirmed via confocal microscopy. In-vivo electrophysiology combined with optical stimulation was used to test optical activation and suppression of MNTB activity during sound stimuli.

**Results:** Opsin expression was strong and stable in MNTB neurons over weeks and months. Laser stimulation during in-vivo recordings successfully induced both activation and suppression of MNTB neurons, demonstrating fast and precise control over neural activity.

**Conclusion:** This in-vivo optogenetic method provides specific, reversible control of MNTB activity in gerbils with rapid, real-time modulation. It offers a powerful tool for investigating sound localization and auditory processing with high temporal precision.

## INTRODUCTION

The medial nucleus of the trapezoid body (MNTB) is a nucleus in the auditory brainstem which provides glycinergic inputs to principal sound localization nuclei, the lateral and medial superior olive (LSO and MSO, respectively) [1–3]. It consists of about 3000-5000 principal neurons [4, 5] which receive contralateral glutamatergic inputs from the globular bushy cells (GBC) of the cochlear nucleus (CN) via a very large synapse known as the calyx of Held [6–8]. MNTB neurons then send glycinergic projections to LSO and MSO [1, 2, 9–14]. These projections are responsible for fast, well-timed inhibitory inputs to MSO and LSO and contribute to computation of interaural time differences (ITD) for low frequency sounds and interaural level differences (ILD) for high frequency sounds, respectively [3, 15–18]. These ITDs and ILDs are critical for allowing listeners to accurately localize sound sources of varying frequencies in the azimuthal plane. Experimental manipulations of the MNTB, therefore, have the potential to offer detailed insights into the mechanisms of sound localization in mammals. In the past, several methods have been used for such manipulations, including ablation [19], pharmacological manipulations [16, 20], and genetic mutation [21]. All of these methods are either irreversible or can only be reversed over a prolonged period. Additionally, these methods are associated with relatively low specificity.

Optogenetics is a powerful alternative method which promises quick reversibility and high specificity of manipulation. While this technique is very well established and heavily used in neuroscience, not much work has been done in auditory brain stem nuclei such as the MNTB. Currently, in-vivo optogenetics in the auditory system has been particularly focused on hearing restoration, with studies targeting the cochlea[22, 23], cochlear nuclei[24, 25], and inferior colliculus[26, 27], while the MNTB has been manipulated during brain slice studies[27]. However, there is still a significant gap in the exploration of optogenetic interventions in this critical auditory region. In the auditory brainstem, several obstacles must be addressed in order to successfully achieve optogenetic control of the target nuclei. First, MNTB is a very deep brain nucleus, roughly 8.5mm from the surface of the brain in the gerbil model [28], making accurate stereotactic targeting for viral injection and electrode insertion challenging. Additionally, this brain area is highly myelinated [3], resulting in optically dense tissue with very high scattering coefficients [29]. Finally, the use of the Mongolian gerbil (*Meriones unguiculatus*), a model system in which very little optogenetics work has been done, requires additional steps to validate optogenetic approaches. There is no transgenic gerbil to date, and the gerbil genome has only been sequenced relatively recently [30–32]. For this reason, transgenic gerbils expressing the required opsins are not available. Instead, viral vectors had to be injected to express these proteins, requiring a significant amount of preliminary testing and validation to determine which viral constructs yield acceptable levels of opsin expression.

Mongolian gerbils are widely employed as a model organism for auditory system research [33]. Unlike mice and rats, gerbils possess an audiogram that covers a significant portion of the human low-frequency hearing range, making them an ideal animal model for investigating sound localization, especially low frequency sound localization, and associated hearing disorders [34, 35]. In the present study, we demonstrate a method for optogenetic manipulation of the MNTB in Mongolian gerbils. To this end, we sterotactically injected MNTB with a variety of optogenetic viruses and quantified opsin expression using confocal microscopy. We then conducted in-vivo optogenetic experiments to validate the optogenetic effects of the expressed opsins.

The method described here has potential to be used in a number of experiments aimed at probing the effects of afferent MNTB inhibition on downstream nuclei such as MSO and LSO. This method provides a powerful tool for probing circuit functionality in the auditory brainstem and exploring the interactions between its nuclei.

## METHODS

### Animals

All experiments were performed on young-adult Mongolian gerbils (*Meriones unguiculatus*). Gerbils were initially purchased from Charles River Laboratories (Wilmington, MA) then bred in-house. All experimental procedures were conducted in compliance with relevant laws, regulations and OLAW guidelines provided by the National Institute of Health and were approved by the University of Colorado Anschutz Medical Campus Institutional Animal Care and Use Committee.

The overall strategy was to stereotactically inject wild type gerbils with adeno-associated viral constructs (AAVs) to express optogenetic proteins. This approach was necessary since no transgenic gerbil lines expressing optogenetic proteins exist to date. The sequence of procedures started with a craniotomy and injections of viral constructs into the MNTB of gerbils, followed by recovery and an incubation period of at least three weeks to allow for sufficient expression of the transgene. After the incubation period, we performed a second procedure in which the optrode was inserted either into MNTB or MSO, and extracellular recordings were performed while the laser was activated during certain periods of the experiment. The overall experimental design is shown in Fig. 1.

**Fig. 1.**
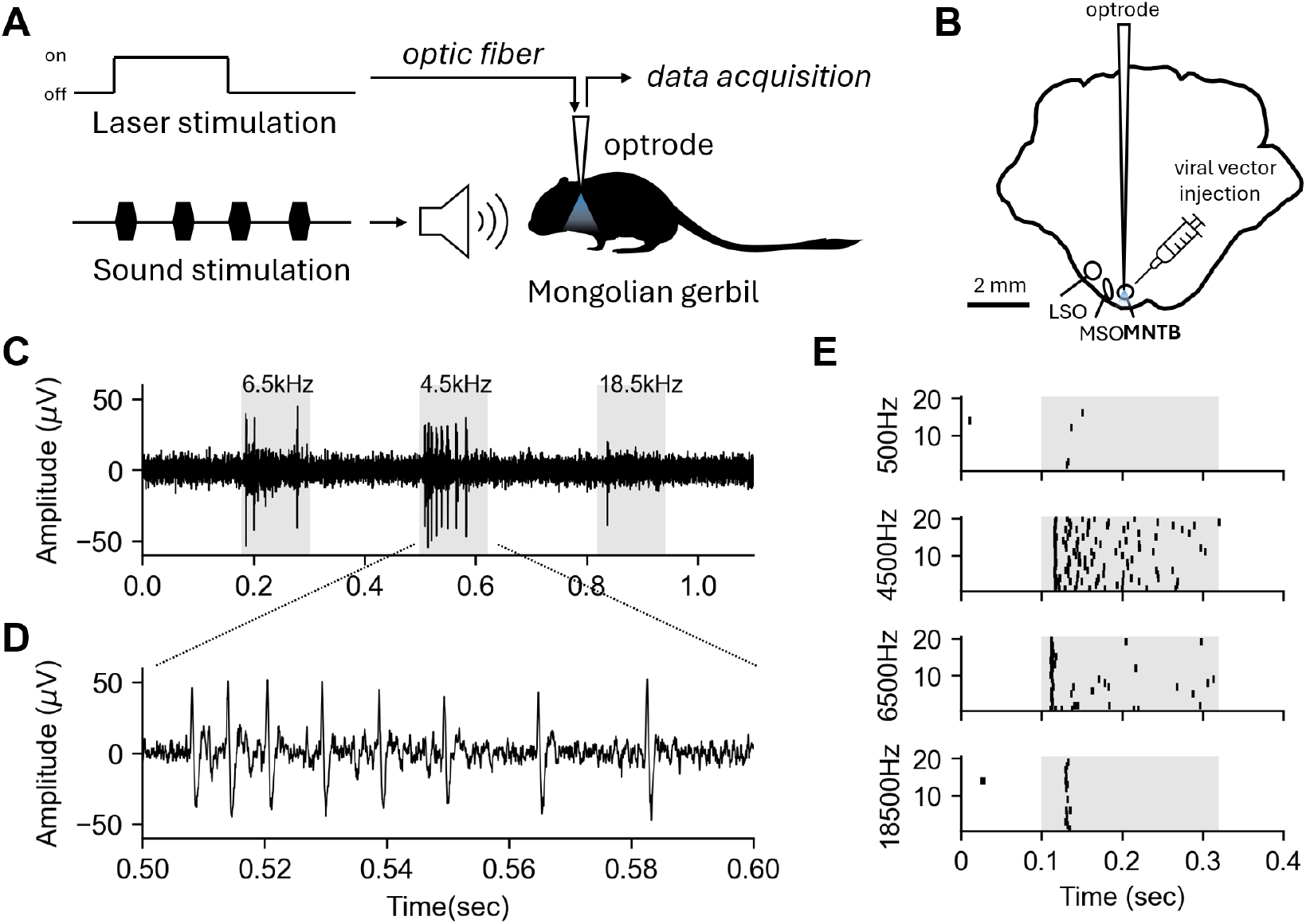
Schematic of experimental design. **A** Diagram showing sound stimulation in the presence and absence of laser stimulation. **B** Location of the viral injection and the recoding sites in the brainstem. **C-E** Example data from electrophysiology recording under auditory stimulation with different sound frequencies. Grey area signifies sound presentation.

### Viral injection

Gerbils were initially anesthetized via isoflurane inhalation (Fluriso, VetOne, ID, USA) in an induction box (5% isoflurane or less). Anesthesia was subsequently maintained in 3% isoflurane or less, guided by lack of the toe-pinch reflex. Animals were prepared for surgery by shaving the skin of the head and applying Betadine and 70% ethanol to the surgical area. The skin overlaying the skull was reflected, and the muscle at the base of the skull was pushed back to uncover the craniotomy site. Stereotaxic coordinates for the MNTB were derived from a gerbil brain atlas [28]. Using a dental drill, a hole was drilled 4mm caudal and 0.8mm lateral to Lambda. A glass pipette was inserted at an angle of 20 degrees relative to the caudal aspect of the skull and the electrode was advanced to a depth of 7500 µm. Once at the target location, 483 nL of optogenetic virus (Table.1) were injected into the MNTB using a nanoliter injector (WPI, Sarasota, FL) (Fig. 1B). The total injection volume was divided into 15 injections of 32.2 nL each, which were injected at a rate of one injection per 15 seconds to achieve the total volume of 483 nL of virus over about 4 minutes. The pipette was retracted, and surgical wounds were sutured using dissolvable polyglactin sutures (Veter Sut, Fort Myers, FL, USA). At the conclusion of the procedure, gerbils were injected subcutaneously with 1mg/kg buprenorphine SR (Wedgewood Connect, San Jose, CA, USA) for post-surgical analgesia. Animals were allowed to recover, and the virus was allowed to incubate for at least three weeks to allow for the expression of opsins before any further procedures were performed.

**Table 1.**
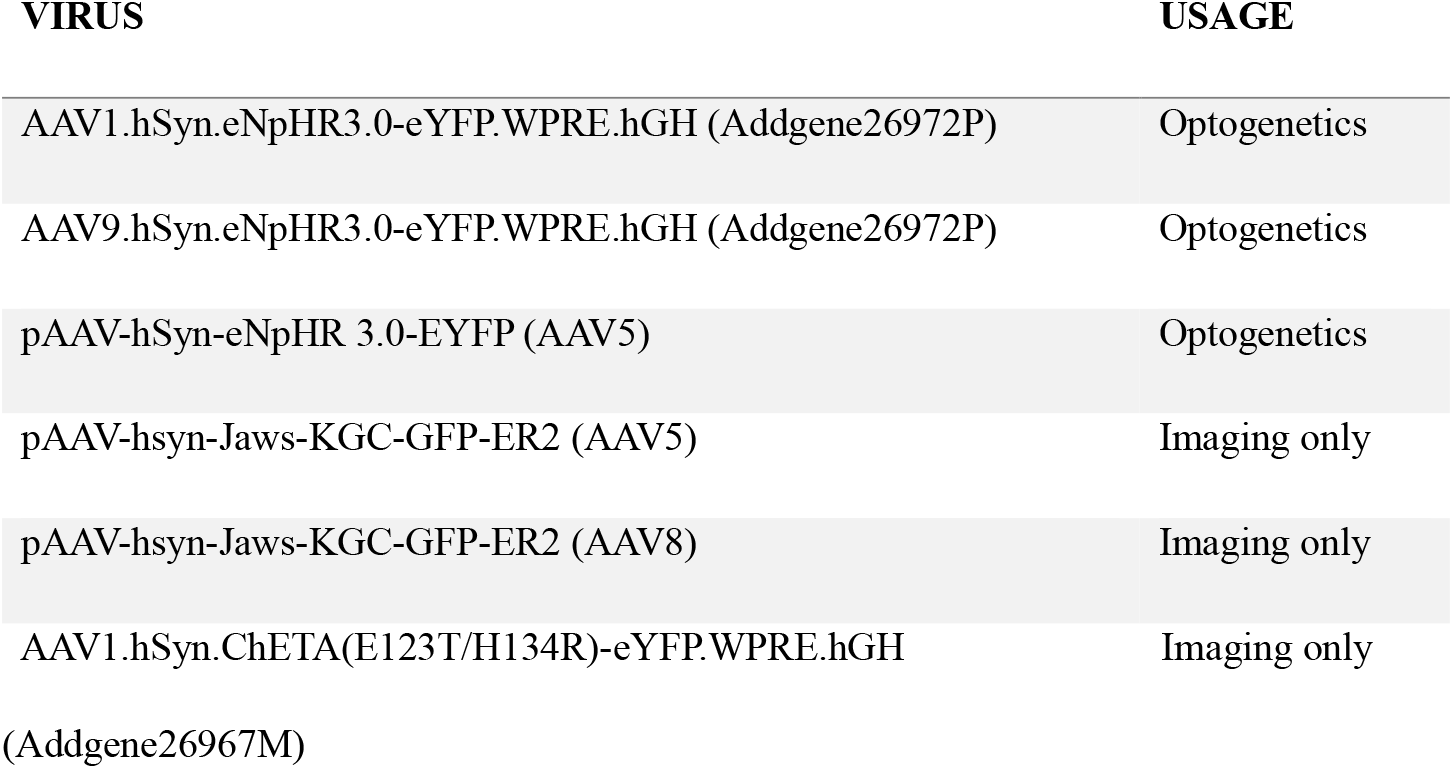
List of injected viral vectors.

| **VIRUS** | **USAGE** |
| --- | --- |
| AAV1.hSyn.eNpHR3.0-eYFP.WPRE.hGH (Addgene26972P) | Optogenetics |
| AAV9.hSyn.eNpHR3.0-eYFP.WPRE.hGH (Addgene26972P) | Optogenetics |
| pAAV-hSyn-eNpHR 3.0-EYFP (AAV5) | Optogenetics |
| pAAV-hsyn-Jaws-KGC-GFP-ER2 (AAV5) | Imaging only |
| pAAV-hsyn-Jaws-KGC-GFP-ER2 (AAV8) | Imaging only |
| AAV1.hSyn.ChETA(E123T/H134R)-eYFP.WPRE.hGH<br>(Addgene26967M) | Imaging only |

### Optrode fabrication

A homemade optrode consisting of a 4 megohm tungsten electrode (WE(25mm)PT34.0B10, Microprobes for Life Sciences, Gaithersburg, MD, USA) and an optical fiber with a core diameter of 105 µm (FG105UCA, Thorlabs, Newton, NJ, USA) was constructed using methods described previously [36] (Fig. 2). For an in-depth optrode fabrication protocol, see protocols.io. Briefly, cannulas were first constructed by stripping 105 µm core optical fiber using a fiber stripper (T06S13), Thorlabs, Newton, NJ). The stripped fiber was cleaved using a fiber cleaver (FC-6S, Sumitomo Electric, Osaka, Japan) and inserted into a 1.25mm ceramic ferrule (MM-FER2007C-1270, Precision Fiber Products, Chula Vista, CA, USA) such that 19mm of fiber protruded from the concave end of the ferrule. To fix the stripped fiber in the ferrule, a drop of 5-minute epoxy (Tank Bond, Dap Products, Baltimore, MD, USA) was added to the concave end of the ferrule before the fiber was inserted, and the fiber was pushed through the epoxy. The fiber was scored and broken off flush with the convex end of the ferrule using a fiber scribe (S90C, Thorlabs, Newton, NJ, USA). Cannulas were then joined to the 25mm tungsten electrodes under a microscope (LaborLux11, Leitz, Stuttgart, Germany). To accomplish this, the rubber insulating sleeve was cut from the electrode’s gold pin connector with a razor blade, and the pin was inserted into a pin header connector held in place under the microscope lens by custom made clip attached to a microscope slide. The cannula was placed on a manipulator stage (423 series, Newport Corporation, Irvine, CA, USA) and the electrode and the cannula were aligned in the same plane of focus (Fig. 2), with the tip of the fiber touching the electrode, and with the electrode tip protruding by 100 µm, as measured by an eyepiece micrometer. UV cure dental cement (NV11VXA, Pentron, Orange, CA) was carefully applied with a scrap piece of optical fiber and cured. Additional dental cement was added and cured as necessary to cover the entire optical fiber, keeping the fiber and the electrode as close together as possible. During this process, some tension was applied to the optical fiber such that the fiber and tungsten electrode were running side by side as parallel as possible, rather than forming a “wedge”“. This was done to keep the optrode diameter as small as possible to minimize brain damage during the insertion. Completed optrodes were connected to the laser via patch cable (RWD Life Science, San Diego, CA) using index matching gel (Thorlabs, Newton, NJ) and ceramic split sleeves (Precision Fiber Products, Chula Vista, CA).

**Fig. 2.**
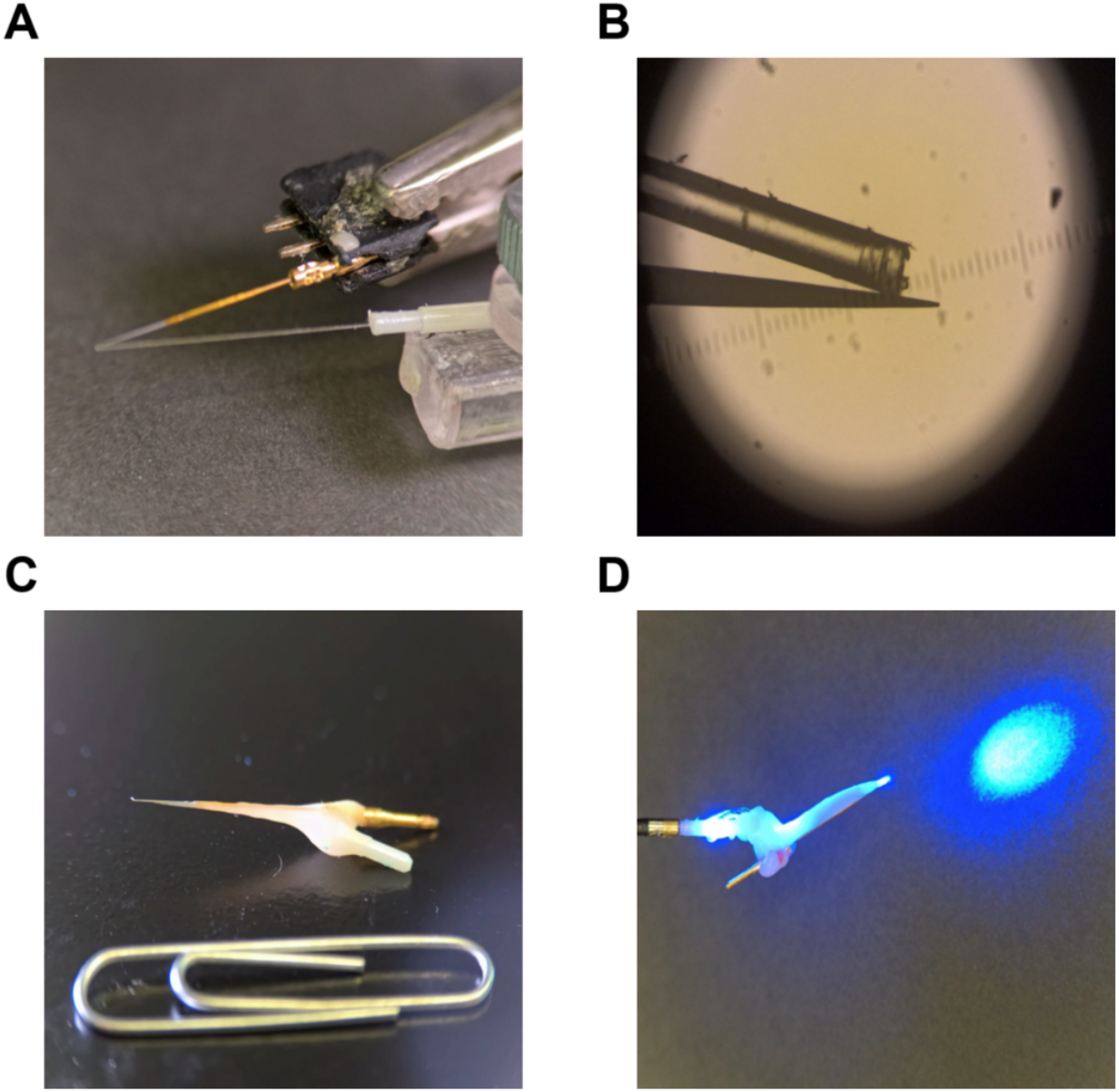
Optrode fabrication. **A-B** Alignment of the tungsten electrode with the glass fiber under a microscope. The completed cannula is maneuvered with manipulator state such that the electrode tip protrudes beyond the fiber tip by 100 µm. **C** A complete optrode with a paperclip for scale. **D** An optrode connected to a blue laser via a patch cable, ceramic sleeve, and index matching gel.

### In-vivo recordings with optogenetic manipulations

Gerbils were initially anesthetized with isoflurane and a craniotomy was performed with the same method as described for the viral injection. Upon the completion of the craniotomy, isoflurane administration was ceased and animals were continuously anesthetized with intraperitoneal injections of a ketamine/xylazine mixture (induction dose = 5mg ketamine/ 100g BW and 0.25mg per 100g BW xylazine, maintenance doses = 1.5mg - 3mg ketamine/ 100g BW approximately every 20-30 minutes according to depth of anesthesia) and placed on a stereotaxic platform in an anechoic chamber with the configuration illustrated in the schematic (Fig .1A). An optrode was inserted and advanced into the brain towards the MNTB with a micromanipulator (IVM1000, Scientifica, Uckfield, UK) (Fig. 1B) using the same stereotaxic coordinates described above. A ground reference was placed by using a needled electrode inserted into the muscle of the dorsal neck. Extracellular neural signals captured by the optrode were pre-amplified using a custom-made bio-amplifier board and sampled at a 50kHz sampling rate using a NIDAQ (National Instrument, TX, USA). Optical stimulation was accomplished using lasers with wavelengths of 635 nm for halorhodopsin- type opsins (Opto Engine LLC, Midvale, UT, USA) and 473 nm for channelrhodopsin-type opsins (SLOC lasers, Shanghai Laser & Optics Century, Shanghai, China). Pure-tone auditory stimuli with different frequencies and intensities were delivered by a pair of multi-field magnetic speakers (MF1, Tucker-Davis Technologies, FL). A recording was recognized as an MNTB neuron if it exhibited tonic or primary like auditory-evoked responses from the contralateral side only. After data acquisition, single-unit spike trains were isolated by performing semi-automatic spike sorting using Gaussian-mixture clustering on bandpass-filtered traces (300-5kHz, 4^th^ order Butterworth) (Fig. 1E). The sorted spike trains were segmented by trials and re-arranged based on the sound stimulus timing and laser condition to generate raster plots and peristimulus time histograms (PSTH).

### Histology

After recordings were complete, gerbils were euthanized with an overdose of pentobarbital (Fatal-Plus, Vortech Pharmaceuticals, Dearborn, MI, USA) and transcardially perfused with phosphate buffered saline (PBS) followed by 4% paraformaldehyde in PBS. Brains were removed and post-fixed overnight in 4% paraformaldehyde, followed by 3 × 10-minute washes in PBS. 100 µm slices were cut with a vibratome (Leica VT 1000 s, Nussloch, Germany), stained with NeuroTrace nissl stain, (ThermoFisher, Waltham, MA, USA) and imaged with a confocal microscope (FV1000, Olympus, Tokyo, Japan) to confirm the presence of EYFP or GFP indicating successful viral expression in MNTB and recording locations. All viruses were imaged to assess expression, but only eNpHR3.0 viruses were used in optogenetic experiments (Fig. 4).

The extent of the virus expression was quantified using standard image processing approaches. Briefly, contours of cells were detected in the Nissl channel using the Sobel filter and Otsu thresholding algorithms [37]. Optogenetic protein expression on the membrane surface was calculated as the summed intensity of the channel expression on the expanded areas of the detected contours. The calculated expressions were normalized to the range between the maximum intensity and average intensity over the entire coronal section to reduce the effects from exposure and autofluorescence. The radius of expression was assessed by computing the half-width of the spatially convoluted expression regions on the membrane. The image processing was implemented using functions from the scikit-image library [38].

## RESULTS

The overarching goal of this project was to determine how optogenetic approaches could be used in a nonstandard animal model, the Mongolian gerbil. While many optogenetic constructs have been developed and tested in mice and rats, very little optogenetic work has been done in gerbils, and the efficacy of these constructs has yet to be tested. Furthermore, there has been limited investigation into optogenetic manipulations of the brain stem, which, again, required different virus envelope types and promoters. Additionally, auditory brain stem nuclei are located near the ventral border of the brain stem, which presents its own issues. For experimental reasons, the surgical and stereotaxic approach of electrodes, optrodes and other tools must be done from the dorsal surface. This is because the middle ear of the gerbil is very large, precluding an approach from the side of the skull. A ventral approach would require a very invasive surgical procedure in which the jaw must be removed [39], which would preclude the required three-week survival of the animals to allow for virus expression. Finally, the target nuclei are located in an excessively myelinated brain area. Myelin increases the scattering coefficients of brain tissue substantially, so a greater light intensity is required to ensure light diffusion and activation of optogenetic proteins in the target area. With these four requirements in mind, we attempted to develop a set of modified techniques that would allow for optogenetic manipulation of the auditory brain stem in Mongolian gerbils.

Various viral constructs with different promotors were tested. We found that constructs (Table. 1) utilizing a human synapsin promotor (SYN) caused spatially concentrated expression of optogenetic proteins in MNTB. By contrast, forebrain-based promotors, such as CaMKII, did not cause any expression of the transgene in MNTB (data not shown). Constructs under the control of SYN and expressing either JAWS or eNpHR 3.0 were stereotactically injected into the MNTB. Confocal microscopy of the entire coronal section and the quantification of membrane expression (Fig. 3) indicates that the viral expression is centered at the MNTB with a diffusion range below 1 mm and radius of less than 500 µm. To assess the time course of the viral expression, confocal microscopy images of the MNTB region were taken at various time points post-viral vector injection (Fig. 4 and Fig. 5). These images clearly show the expression of injected viral constructs on the cell membrane of MNTB principal neurons after 2-3 weeks of incubation. Moreover, the expression is stable over months and still salient after 200 days post injection, suggesting that optogenetic manipulation of MNTB is possible for long-term experiments.

**Fig. 3.**
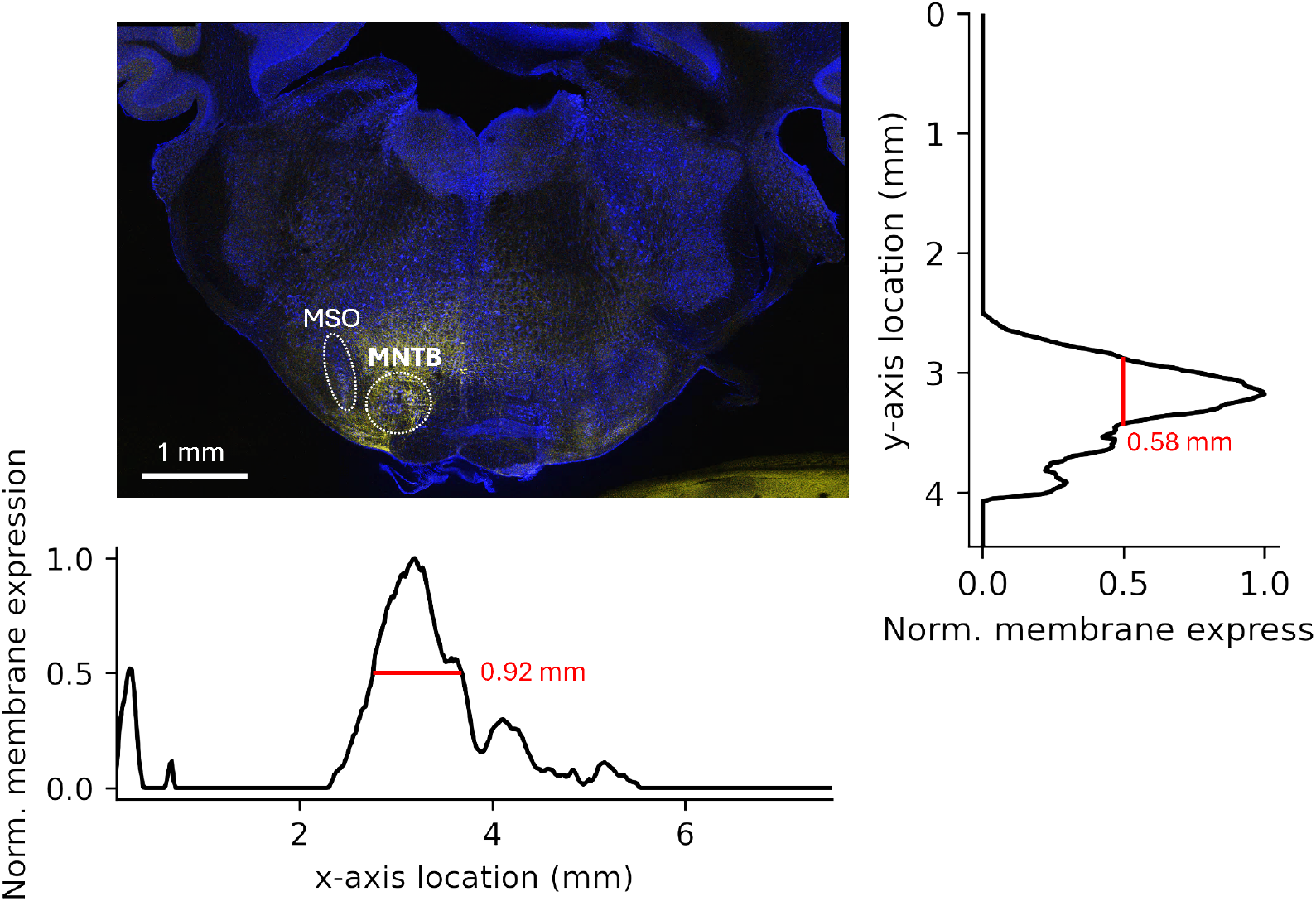
Confocal image of opsin expression in an auditory brainstem section and quantification for spatial diffusion of viral expression. (blue: Nissl; yellow: opsin; scale bar: 1 mm) Expression is well localized to MNTB.

**Fig. 4.**
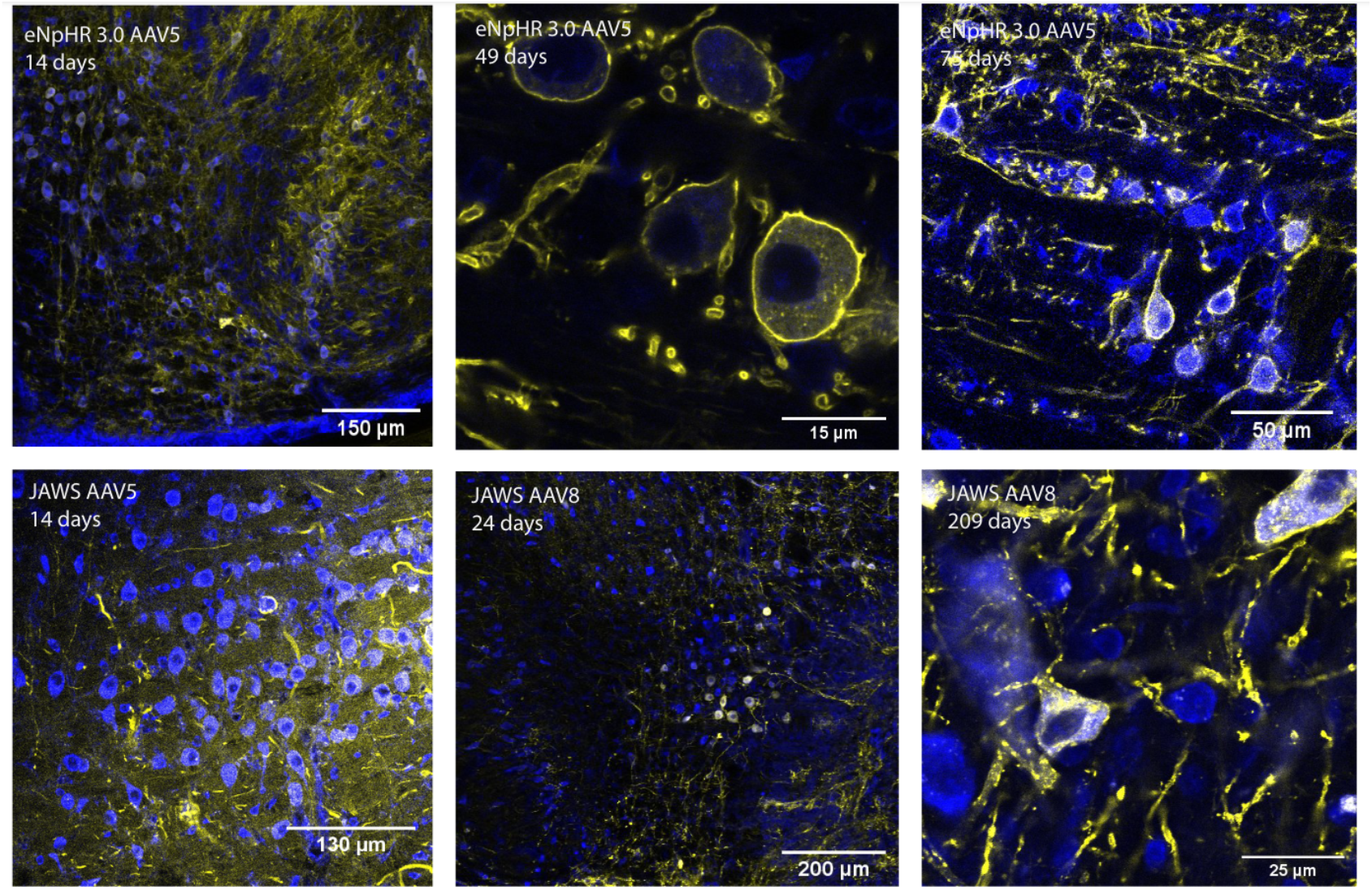
Confocal images of eNpHR 3.0 AAV5 expression and JAWS AAV8 expression over time after viral injection. Both viral constructs yielded obvious, long-lasting expression of viral proteins (blue: Nissl; yellow: opsin). Top row: eNpHR 3.0 AAV5 Expression over time (14 days, 49 days, 75 days). Bottom Row: JAWS AAV8 expression over time (14 days, 24 days, 209 days).

**Fig. 5.**
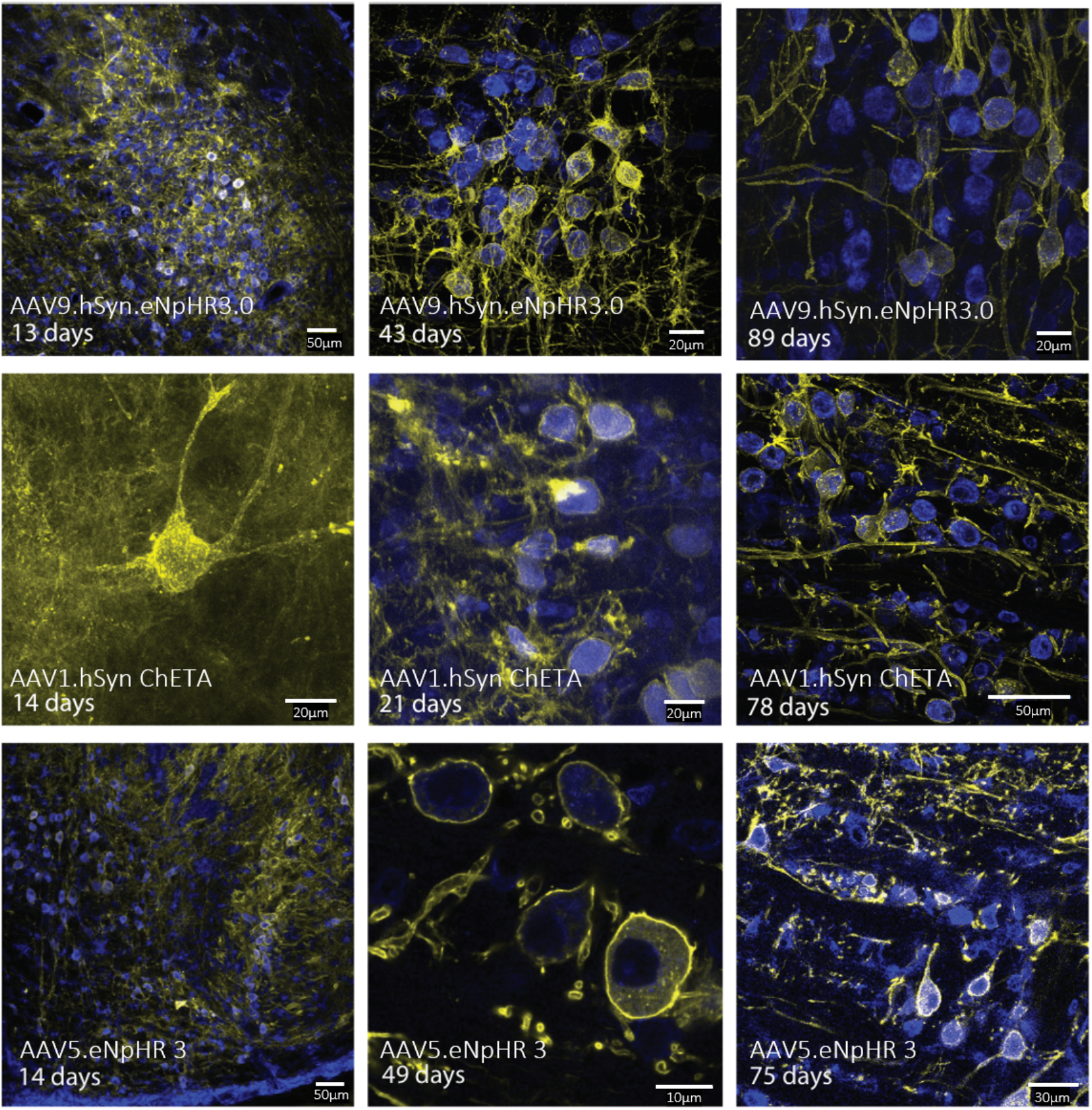
Confocal images of MNTB sections from gerbils injected with optogenetic viral constructs to test expression levels and longevity of the expression. Top row: AAV9.hSyn.eNpHR3.0; center row: AAV1.hSyn ChETA; Bottom row: AAV5.eNpHR 3. (blue: Nissl; yellow: opsin).

In-vivo optogenetic experiments were performed on anesthetized gerbils along with the presence of optical and auditory stimulation (Fig. 1A). The light-evoked suppression was observed in MNTB neurons recorded from Halorhodopsin-injected gerbils as seen in the example shown in Fig. 6. When the red-color laser was presented, the auditory tuning characteristics of the MNTB neuron changed systematically with a reduction of the sound evoked firing rates by half (Fig. 6A). Much less spontaneous activity before and after sound stimuli was also observed in laser-on trials (Fig. 6B, C). Using the same approach on gerbils injected with channelrhodopsin-type viral vector, the light-evoked excitation was shown in recorded MNTB neurons as seen in Fig. 7. The ChR-2 expressing MNTB neurons fire twice as many spikes in response to sound stimuli while under blue-laser stimulation (Fig. 7A). Spontaneous activities are also substantially increased under blue-laser stimulation (Fig. 7B, C). These experiments demonstrate that the in-vivo neural activities of MNTB cells can be effectively manipulated using optogenetic approaches.

**Fig. 6.**
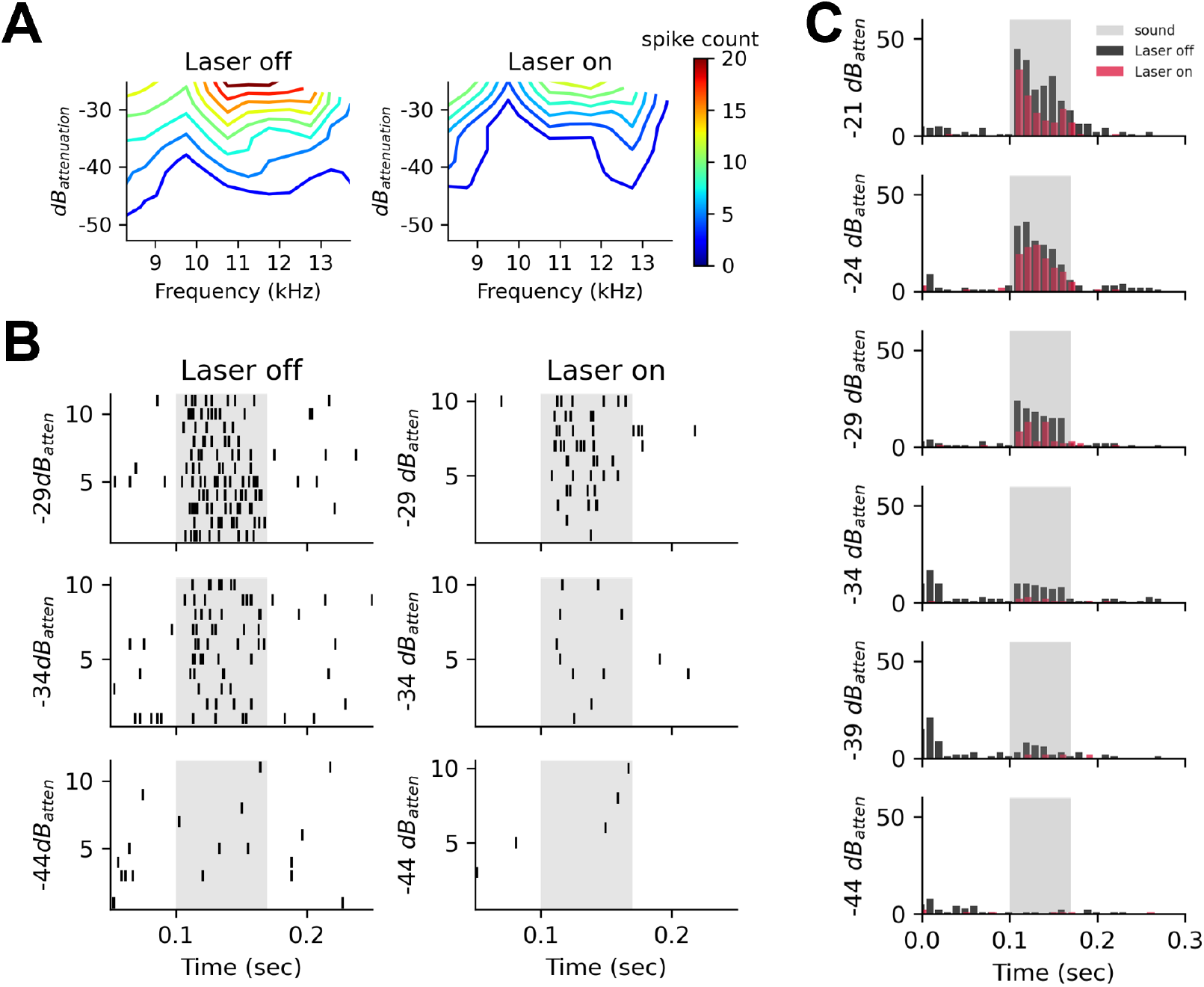
Optogenetic manipulation of a Halorhodopsin-expressing MNTB cell. **A** Cell firing-rate tuning under different sound levels and frequencies with and without laser stimulation. **B** Raster plots of MNTB cell over 10- trial replicates. Gray shades indicate auditory stimulus window. **C** PSTH of the MNTB cell over 10-trial replicates under various laser conditions. Gray shades indicate the auditory stimulus window.

**Fig. 7.**
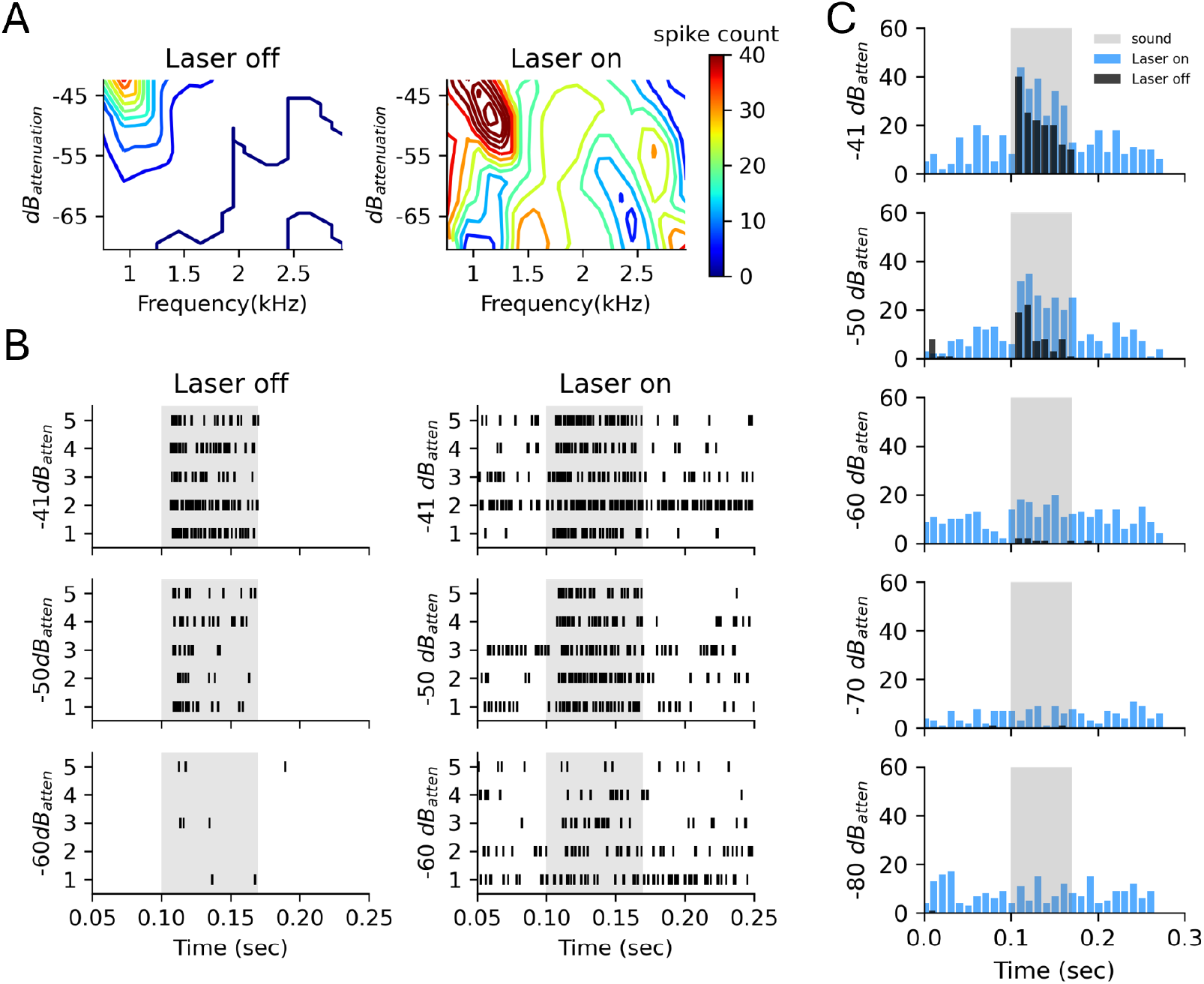
Optogenetic manipulation of a ChR2-expressing MNTB cell. **A** Cell firing-rate tuning under different sound levels and frequencies with and without laser stimulation. **B** Raster plots of the MNTB cell over 10-trial replicates. Gray shades indicate the auditory stimulus window **C** PSTH of the MNTB cell over 10-trial replicates under various laser conditions. Gray shades indicate the auditory stimulus window.

To validate the effectiveness of the proposed optogenetic manipulation, we performed an electrophysiological experiment on an age-matched wild-type gerbil which did not receive any viral injection. This was done to test for potential optical and heating effects of laser stimulation. No specific light-induced effects were found in the MNTB neurons. Fig. 8 shows recordings from a single gerbil. The presence of the laser at maximum power had no impact on the noise floor (Fig. 8A), firing activity, or temporal dynamics from PSTHs (Fig. 8B-E). These results suggest that the optogenetic manipulations described are specific to opsin presence and are not driven or influenced by the light or heating effects from the laser.

**Fig. 8.**
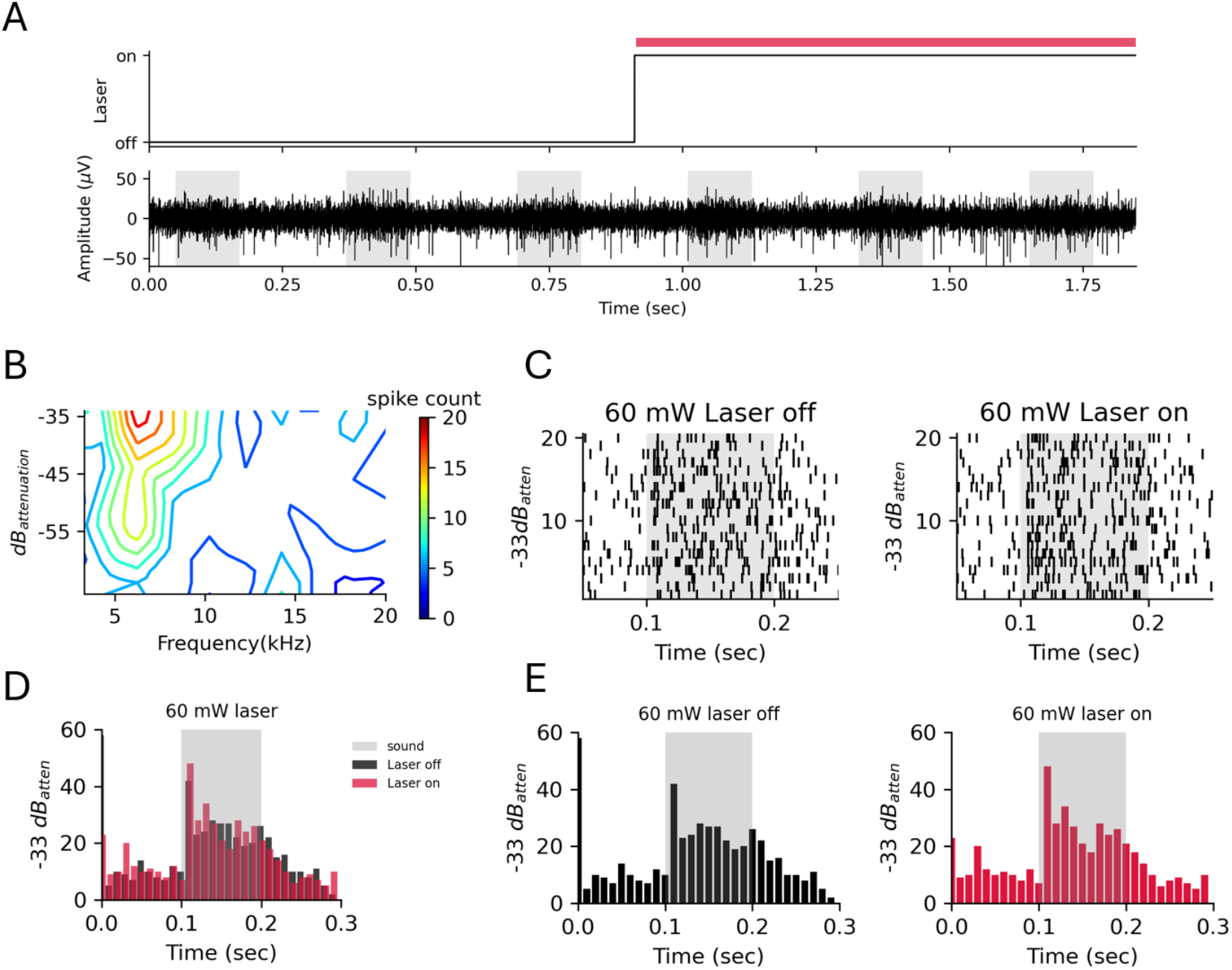
Electrophysiological recordings from the MNTB of a wild-type gerbil which did not receive any viral injection. **A** Raw recording trace (red bar: laser onset period; gray shade: sound stimulation). **B** Cell firing rate tuning under different sound levels and frequencies. **C** Raster plots of the MNTB cell firing over 10-trial replicates. Gray shades indicate auditory stimulus window. **D-E** PSTH of the MNTB cell over 10- trial replicates under various laser conditions. Gray shades indicate the auditory stimulus window.

## DISCUSSION

In this study, we demonstrate an approach to in-vivo optogenetic manipulation of gerbil MNTB neurons. We first confirm the expression of various optogenetic viruses containing a human synapsin promoter and either a GFP or EYFP fusion protein in the gerbil MNTB region. Both JAWS and eNpHR 3.0 opsins, when delivered via adeno-associated viruses, were highly expressed in MNTB principal neurons. Furthermore, opsins remain present for extended periods of time after injection, allowing for effective control of the target region in both short-term and long term-experiments. Finally, we used in-vivo optogenetic experiments to modulate MNTB activity via laser stimulation.

These results validate the proposed approach and enable in-vivo experiments in which MNTB activity can be manipulated in more complex ways. Such experiments allow for the investigation of functional roles of MNTB derived inhibition in the afferent auditory pathway and the sound localization process. MNTB outputs are inhibitory projections to multiple other nuclei of the auditory brainstem [1, 40–43] and play a critical role in the localization of sound, so experimental alteration of its firing allows for more in-depth studies of these processes. For example, the medial and lateral superior olivary nuclei (MSO and LSO) both receive inhibitory inputs from MNTB, and their neural processing can be studied by altering these MNTB inhibitory inputs [3]. Several studies manipulated the inhibitory inputs to these nuclei using pharmacological approaches with strychnine iontophoresis. When MSO neurons were manipulated with this approach, ITD tuning characteristics were potentially influenced [16, 20, 44]. Furthermore, a more recent computational model predicts that timing and strength of the MNTB input shapes ITD tuning curves and contributes to the temporal precision of ITD detection [45]. However, these findings have the limitation that pharmacological manipulation happens at the target site, disallowing the investigator to make a final determination about the anatomical source of the inhibition. Additionally, iontophoresis is associated with slow time constants, making it difficult to control diffusion dynamics, concentration dependent effects [46], and complete blockage of all inhibition is atypical for most in-vivo situations. Optogenetic manipulations have the potential to overcome these drawbacks and may facilitate the design of more sophisticated experiments to address the shortcomings of modern methods.

Optogenetic manipulations may also be ideal for studying other deep auditory brain stem nuclei which are poorly understood. For example, the lateral nucleus of the trapezoid body (LNTB) is believed to send inhibitory projections to its ipsilateral MSO [47], but the functional role of this projection is unclear. Similar methods to those described here could be used to explore the role of this nucleus in sound localization. Furthermore, several deficits in spatial hearing, including those related to Fragile X Syndrome (the most common monogenic form of autism) are postulated to be related to pathologies in this system [48]. Clearly, the ability to experimentally manipulate this system offers great potential to test pathological hypotheses and eventually suggest insights into potential treatments.

## CONCLUSION

Optogenetics is a powerful, widely used tool for the manipulation of neural circuits, but its use in the auditory brainstem has been very limited. Here, we demonstrate the usefulness of this technique in MNTB and validate the use of several viruses in the superior olivary complex (SOC). Additionally, we present a method to successfully overcome the obstacles inherent to in-vivo optogenetic manipulations of the auditory brainstem. Namely, the small, deep nuclei of the SOC can be accurately and precisely targeted for stereotactic injection of optogenetic viruses, and laser stimulation can be delivered concurrently with sound stimulation using custom optrodes such as those described here. Mongolian gerbils are frequently used in auditory research due to their comparable hearing range to humans. Demonstrating the effectiveness of various viral constructs in their auditory brainstem is a key step in investigating questions related to the functionality of these circuits in the human auditory brainstem. We demonstrate both excitatory and inhibitory manipulations as well as provide evidence for the effective long-term infection of MNTB neurons with several optogenetic viruses. Our results provide evidence for potential use of this method across a variety of experimental designs and timelines.

## Acknowledgement

We would like to thank Ajay Keerthy for helpful discussions and critically reading the manuscript. We would like to thank the personnel from the University of Colorado Anschutz Medical Campus Advanced Light Microscopy Core for imaging assistance. Imaging was performed in the Advanced Light Microscopy Core facility of the NeuroTechnology Center at the University of Colorado Anschutz Medical Campus, which is supported in part by Rocky Mountain Neurological Disorders Core Grant (P30NS048154) and by Diabetes Research Center Grant (P30 DK116073).

